# A machine-readable specification for genomics assays

**DOI:** 10.1101/2023.03.17.533215

**Authors:** A. Sina Booeshaghi, Xi Chen, Lior Pachter

**Author notes:** Address correspondence to &.

## Abstract

Understanding the structure of sequenced fragments from genomics libraries is essential for accurate read preprocessing. Currently, different assays and sequencing technologies require custom scripts and programs that do not leverage the common structure of sequence elements present in genomics libraries. We present *seqspec*, a machine-readable specification for libraries produced by genomics assays that facilitates standardization of preprocessing and enables tracking and comparison of genomics assays. The specification and associated *seqspec* command line tool is available at https://github.com/IGVF/seqspec.

## Introduction

The proliferation of genomics assays (Ogbeide et al. 2022) has resulted in a corresponding increase in software for processing the data (Zappia, Phipson, and Oshlack 2018). Frequently, custom scripts must be created and tailored to the specifics of assays, where developers reimplement solutions for common preprocessing tasks such as adapter trimming, barcode identification, error correction, and read alignment (Wu et al. 2022; Ma et al. 2020; Cheow et al. 2016; Healey, Bassham, and Cresko 2022). When software tools are assay specific, parameter choices in these methods can diverge, making it difficult to perform apples-to-apples comparisons of data produced by different assays. Furthermore, the lack of preprocessing standardization makes reanalysis of published data in the context of new data challenging.

While genomics protocols can vary greatly from each other, the libraries they generate share many common elements. Typically, sequenced fragments will contain one or several “technical sequences” such as barcodes and unique molecular identifiers (UMIs), as well as biological sequences that may be aligned to a genome or transcriptome. Standard library preparation kits generally require that DNA from the libraries is cut, repaired, and ligated to sequencing adapters (Figure 1). Primers bind to the sequencing adapters, and initiate DNA sequencing whereby reads are subsequently generated. Illumina sequencing employs a sequencing by synthesis approach where fluorescently labeled nucleotides are incorporated into single-stranded DNA, and imaged, while PacBio uses zero-mode waveguides for single-molecule detection of dNTP incorporation. Oxford Nanopore on the other hand binds sequencing adapters to pores in a flow cell and DNA is sequenced by changes in electrical resistance across the pore (Iizuka, Yamazaki, and Uemura 2022).

**Figure 1:**
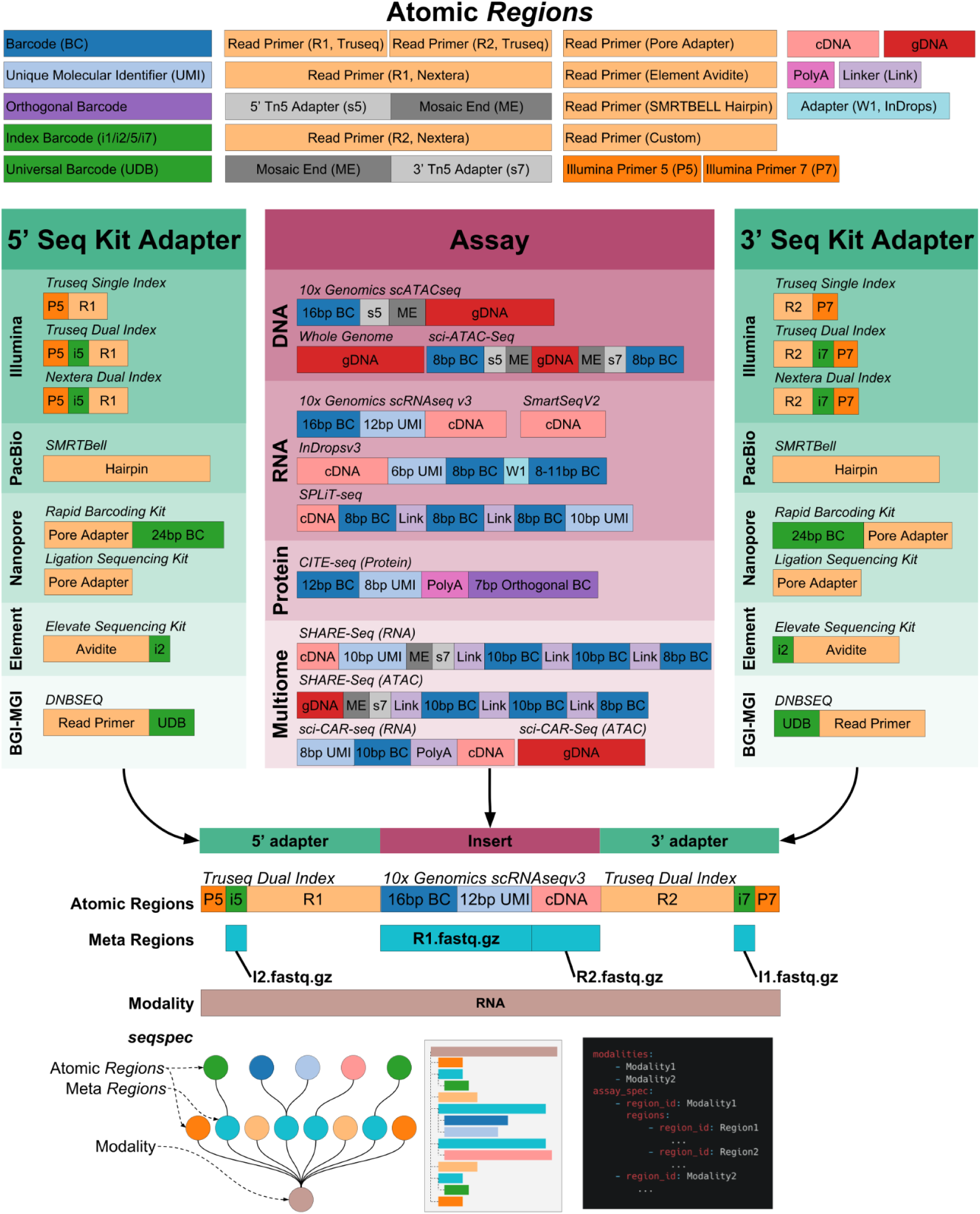
The structure of reads sequenced from genomics libraries. Sequencing libraries are constructed by combining Atomic *Regions* to form an adapter-insert-adapter construct. The *seqspec* for the assay annotates the construct with *Regions* and *Meta Regions*

Many single-cell genomics assays introduce additional library complexity further complicating preprocessing. For example, the inDropsv3 (Klein et al. 2015) assay produces variable length barcodes while the 10x Genomics scRNA-seq assay (Zheng et al. 2017) produces fixed-length barcodes that are derived from a known list of possibilities.

Current file formats such as FASTQ, Genbank, FASTA, and workflow-specific files (Parekh et al. 2018) lack the flexibility to annotate sequenced reads that contain these complex features. In the absence of sequence annotations, processing can be challenging, limiting the reuse of data that is stored in publicly accessible databases such as the Sequence Read Archive (Katz et al. 2022). To facilitate utilization of genomics data, a database of assays along with a description of their associated library structures was assembled in (Chen 2020). While this database has proved to be very useful, the HTML descriptors are not machine readable. Moreover, the lack of a formal specification limits the utility and expandability of the database.

## Results

The *seqspec* specification defines a machine-readable file format, based on YAML, that enables sequence read annotation. Reads are annotated by *Regions* which can be nested and appended to create a *seqspec. Regions* are annotated with a variety of properties that simplify the downstream identification of sequenced elements. The following are a list of properties that can be associated with a *Region*:

- Region ID: unique identifier for the *Region* in the *seqspec*
- Region type: the type of region
- Name: A descriptive name for the *Region*
- Sequence: The specific nucleotide sequence for the *Region*
- Sequence type: The type of sequence (fixed, onlist, random, joined)
- Minimum length: The minimum length of the sequence for the *Region*
- Maximum length: The maximum length of the sequence for the *Region*
- Onlist: The list of permissible sequences from which the Sequence is derived

Importantly, *Regions*, known as meta *Regions*, can contain *Regions*; a property that is useful for grouping and identifying sequence types that are contained in reads. The YAML format is a natural language to represent nested meta-*Regions* in a human-readable fashion. Python-style indentation and syntax can be used to create a human-readable file format without the excessive grouping delimiters of alternative languages such as JSON. Additionally, nested *Regions* allow Assays to be represented as an Ordered Tree where the ordering of subtrees is significant: atomic *Regions* are “glued” together in an order that is concordant with the design of the sequencing library in the 5’ to 3’ direction (Supplementary Figure 1).

Importantly, *seqspec* files are machine-readable, and *Region* data can be parsed, processed, and extracted with the *seqspec* command-line tool. The tool contains six subcommands that enable various tasks such as specification checking, finding, formatting, and indexing,

1. *seqspec check*: check the correctness of attributes against the *seqspec* schema
2. *seqspec find*: print *Region* metadata
3. *seqspec format*: auto populate *Region* metadata for meta *Regions*
4. *seqspec index*: extract the 0-indexed position of *Regions*
5. *seqspec init*: initialize a seqspec with a newick-formatted string
6. *seqspec onlist*: get the path to the onlist file for the specific region type
7. *seqspec print*: print html, markdown, ascii, read diagram that visualizes the *seqspec*
8. *seqspec split*: split a FASTQ file by *Regions*

To illustrate how *seqspec* can be used to facilitate processing and analysis of single-cell RNA-seq reads, we implemented in the *seqspec index* command the facility to produce the relevant technology string for three single-cell RNA-seq preprocessing tools: kallisto bustools (Melsted et al. 2021), simpleaf/alevin-fry (He et al. 2022), and STARsolo (Kaminow, Yunusov, and Dobin 2021) (Figure 2). *Regions* associated with barcodes, UMIs, and cDNA are extracted, positionally indexed and formatted on a per-tool basis. The modularity of *seqspec* makes it simple to produce tool-compatible technology strings for other assay types.

**Figure 2:**
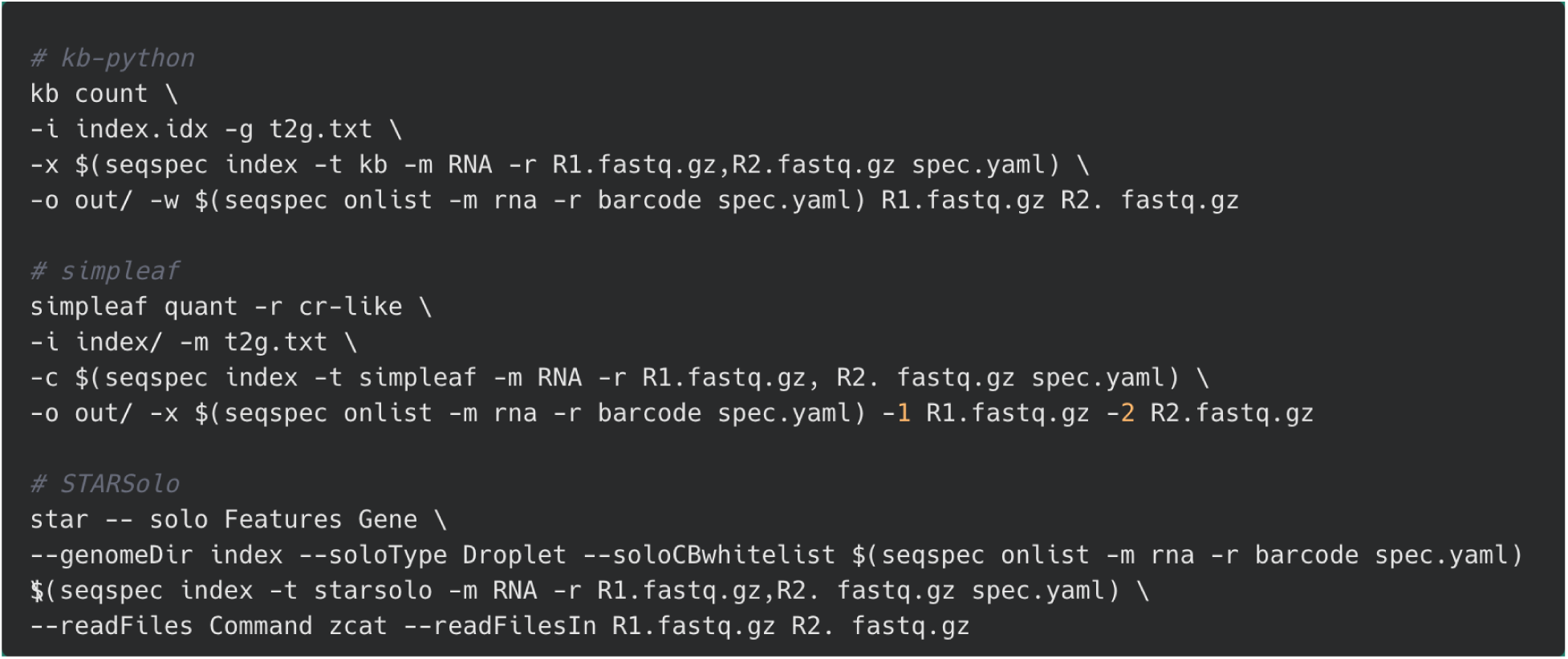
Uniform processing enabled with *seqspec*. The *seqspec index* command produces a technology string that identifies appropriate sequence elements and can be passed into processing tools.

## Discussion

Standardized annotation of sequencing reads in a human- and machine-readable format serves several purposes including the enablement of uniform processing, organization of sequencing assays by constitutive components, and transparency for users. The flexibility of *seqspec* should allow it to be used for all current sequence census assays (Wold and Myers 2008), and specifications should be readily adaptable to different sequencing platforms; our initial release of *seqspec* contains specifications for 38 assays (see https://igvf.github.io/seqspec/).

Comparison of *seqspec*s for different assays, immediately reveals shared similarities and differences that can be visualized with *seqspec print*. For example, the SPLiT-seq single-cell RNA and the multimodal SHARE-seq single-cell assays are aimed at different modalities and utilize different protocols to produce libraries, but the resultant structures are very similar (Figure 1) since they both rely on split-pool barcoding (Rosenberg et al. 2018). The *seqspec* for the sci-CAR-seq assay (Cao et al. 2018), from which split-pool assays such as SHARE-seq are derived, shows that the cell barcoding is encoded in the Illumina indices. It should be possible to develop an ontology of assays by comparing the *seqspec* specifications of assays and quantifying their similarities and differences.

In demonstrating that *seqspec* can be used to define options for preprocessing tools, we have shown that *seqspec* is immediately useful for uniform processing of genomics data. The preprocessing applications will hopefully incentivize data generators to define and deposit *seqspec* files alongside sequencing reads in public archives such as the Sequence Read Archive. While *seqspec* is not a suitable format for general metadata storage, the precise specification of sequence elements present in reads, including sequencer-specific constructs, should be helpful in identifying batch effects even when metadata is missing or inaccurate.

## Acknowledgements

We thank Delaney Sullivan for helpful discussions and Rahma Elsiesy for helpful feedback on Figure 1. Discussions with the Impact of Genomics Variation on Function (IGVF) Single-Cell Focus Group helped to shape some features of *seqspec*. Thanks to Idan Gabdank for useful feedback on *seqspec* and for suggesting the md5 checksum. Meichen Fang contributed the sci-RNA-seq3 *seqspec*. A.S.B. and L.P. were supported in part by NIH 5UM1HG012077-02.

