## Supplementary Information for "A machine-readable specification for genomics assays"

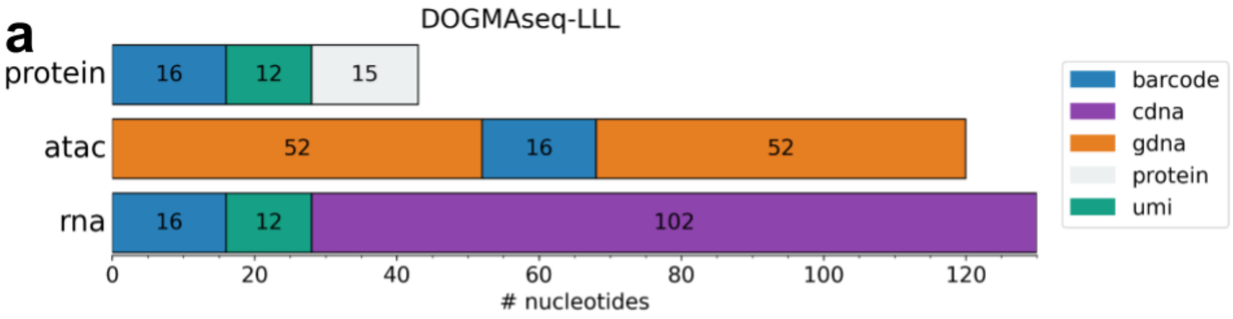


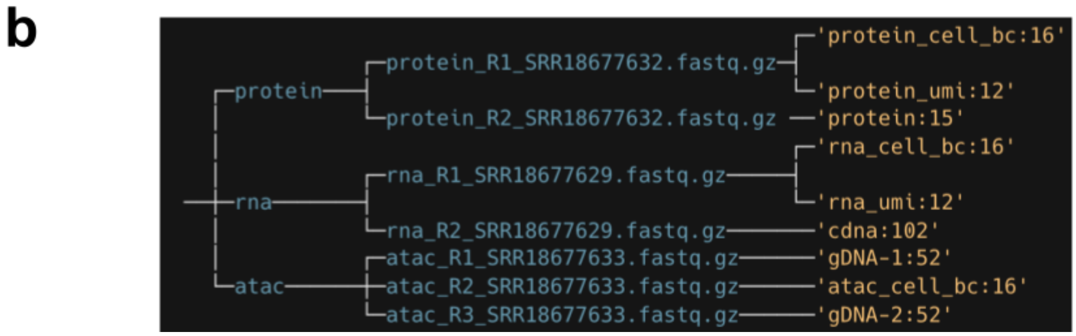


**Supplementary Figure 1:** *seqspec* read structure of the DOGMAseq-LLL (Xu et al. 2022) assay annotated by (a) atomic regions and their lengths. The (b) ordered-tree representation of the reads.
